# Nef enhances HIV-1 replication and infectivity independently of SERINC3 and SERINC5 in CEM T cells

**DOI:** 10.1101/2020.07.15.205757

**Authors:** Peter W. Ramirez, Thomas Vollbrecht, Francisco M. Acosta, Marissa Suarez, Aaron A. Angerstein, Jared Wallace, Ryan M. O’ Connell, John Guatelli

**Affiliations:** Department of Biological Sciences, California State University Long Beach, Long Beach, CA; Department of Medicine, University of California San Diego, La Jolla, California, USA; VA San Diego Healthcare System, San Diego, California, USA; Division of Microbiology and Immunology, Department of Pathology, The University of Utah, Salt Lake City, UT

## Abstract

The lentiviral *nef* gene encodes several discrete activities aimed at co-opting or antagonizing cellular proteins and pathways to defeat host defenses and maintain persistent infection. Primary functions of Nef include downregulation of CD4 and MHC class-I from the cell surface, disruption or mimicry of T-cell receptor signaling, and enhancement of viral infectivity by counteraction of the host antiretroviral proteins SERINC3 and SERINC5. In the absence of Nef, SERINC5 incorporates into virions and inhibits viral fusion with target cells, decreasing infectivity. However, whether Nef’s counteraction of SERINC5 is the cause of its positive influence on viral growth-rate in CD4-positive T cells is unclear. Here, we utilized CRISPR/Cas9 to knockout SERINC3 and SERINC5 in a leukemic CD4-positive T cell line (CEM) that displays relatively robust *nef*-related infectivity and growth-rate phenotypes. As previously reported, viral replication was attenuated in CEM cells infected with HIV-1 lacking Nef (HIV-1ΔNef). This attenuated growth-rate phenotype was observed regardless of whether the coding regions of the *serinc3* or *serinc5* genes were intact. Moreover, knockout of *serinc3* or *serinc5* failed to restore the infectivity of HIV1ΔNef virions produced from infected CEM cells. Taken together, our results corroborate a similar study using another T-lymphoid cell line (MOLT-3) and indicate that the antagonism of SERINC3 and SERINC5 cannot fully explain the virology of HIV-1 lacking Nef.

## Introduction

Primate lentiviruses encode several accessory gene products that facilitate viral reproduction and persistence, in some cases by providing evasion of the host immune response. The lentiviral Nef protein accelerates viral pathogenesis and progression to AIDS in SIV infected rhesus macaques (1) and enhances disease progression in humans infected with HIV-1 (2, 3). Nef is a small, myristoylated, peripheral membrane protein with many well conserved activities, including the downregulation of cell surface proteins such as CD4 and MHC class-I (MHC-I), the modulation of T cell activation, and the enhancement of viral infectivity and growth-rate (4).

To downregulate CD4, Nef binds the cytoplasmic domain of CD4 and links it to the clathrin Adaptor Protein 2 (AP-2) complex, internalizing CD4 from the plasma membrane and delivering it to lysosomes for degradation (5–8). The targeting of CD4 by Nef contributes to viral replication in several ways. CD4 downregulation prevents superinfection of cells and consequent premature cell-death, ensuring adequate time for viral replication (9). It also contributes to inhibiting antibody dependent cellular cytotoxicity (ADCC) that recognizes CD4-induced epitopes with Env (10). Potentially directly relevant to the work presented here, CD4 incorporates into virions and inhibits virion-infectivity if not downregulated by Nef (11, 12).

To downregulate MHC-I, Nef has been proposed to use two non-mutually exclusive mechanisms: 1) Nef and the clathrin-adaptor AP-1 intercept *de novo* synthesized MHC-I molecules within the *trans*-Golgi network (TGN), leading to lysosomal degradation (13– 16) ; and/or 2) Nef retains internalized MHC-I molecules in the TGN via induction of a Src family kinase/phosphoinositide 3-kinase signaling cascade (17–19). In either case, reducing the expression of MHC-I molecules at the plasma membrane protects HIV-1 infected cells from lysis by cytotoxic T-lymphocytes, contributing to immune evasion (20).

Nef interacts directly with Src-family kinases including the lymphocyte specific kinase Lck, which it downregulates from the cell surface (21). The many studies of Nef’s influence on T cell activation are difficult to reconcile, but transcriptional profiling suggests that the expression of Nef mimics signaling through the T cell receptor (22). Apart from the model of MHC-I downregulation noted above, a clear mechanistic link between the influence of HIV-1 Nef on T cell activation and its influence on the expression of cell surface receptors is lacking.

Nef stimulates HIV-1 replication and enhances virion-infectivity in many cell culture systems, including various T cell lines, primary CD4-positive T cells, and human lymphoid tissue (23–26). Diverse *nef* alleles from humans and primates maintain the ability to enhance HIV-1 replication and infectivity, suggesting these functions are important for establishing and maintaining persistent infection (26–28). Expression of Nef within virion-producer cells and encoded either in *cis* or in *trans* relative to the viral genome enhances HIV-1 replication and yields virions of greater infectivity (29, 30). Nef’s ability to enhance infectivity requires cellular components involved in vesicular trafficking (dynamin 2, AP-2, and clathrin) and is also determined by the Envelope (Env) glycoprotein (31, 32). A survey of cell lines identified murine leukemia virus glycosylated Gag (MLV glycoGag), a protein structurally unrelated to Nef, as an infectivity factor that rescues Nef-deficient HIV-1 virions (33). These and other features suggested that Nef counteracts a cellular factor that restricts viral infectivity and possibly replication.

Two groups identified the host transmembrane protein SERINC5, and to a lesser degree SERINC3, as an inhibitor of HIV-1 virion-infectivity that is counteracted by Nef (34, 35). Nef downregulates SERINC5 from the plasma membrane via a clathrin/AP-2 and Rab5/7 endo-lysosomal pathway (34, 36), reducing the incorporation of SERINC5 in HIV-1 virions. This in turn correlates with more efficient fusion of virions with target cells and greater infectivity (34, 35). Nef’s ability to counteract SERINC5 is conserved across primate lentiviruses and correlates with the prevalence of these viruses in the wild (34, 37). Modulation of SERINC5 extends to other retroviral proteins, including S2 from equine infectious anemia virus (EIAV) as well as MLV glycoGag (34, 35, 38). HIV-1 Env glycoproteins are differentially sensitive to SERINC-mediated restriction when produced from CD4-negative cells in single-round replication assays; sensitivity correlates to some extent with the degree of Env-trimer openness and instability (34, 35, 39, 40).

SERINCs comprise a family of five genes that are evolutionarily conserved from yeast to mammals (41). They encode multi-pass transmembrane proteins that support serine specific phospholipid biosynthesis (hence their name: <u>ser</u>ine <u>inc</u>orporator) (42), yet this function does not seem to account for their anti-retroviral activity (43). Rather, SERINC5 appears to disrupt the formation of fusion pores between HIV-1 virions and target cells (44) in an Env-conformation and CD4-dependent manner (45).

Initial studies of the Nef-SERINC relationship focused on the Jurkat T cell line, due to the large defect in the infectivity of Nef-negative virions produced from these cells and their relatively high levels of SERINC5 mRNA. Studies using another CD4-positive T cell line, MOLT-3, have recently cast doubt on whether SERINC family proteins are sufficient to explain the virologic phenotypes of Nef (46). In support of a SERINC-dependent mechanism, Nef does not enhance HIV-1 infectivity and replication-rate in Jurkat T cells when SERINC3 and SERINC5 are knocked out (34, 35). Moreover, a minimal MLV glycoGag (termed glycoMA) can functionally replace Nef with respect to viral replication rate and virion-infectivity when the virus is propagated using Jurkat cells (46). In contrast, Nef, but not glycoMA, enhances HIV-1 replication in MOLT-3 cells. The Nef-effect in MOLT-3 cells persists when the cells are knocked out for SERINC3 and SERINC5, indicating that these restriction factors are not necessary for the virologic effects of Nef in this setting (46). Remarkably, glycoMA cannot substitute functionally for Nef with respect to stimulating viral replication in primary CD4-positive T cells. This suggests that the growth rate enhancing effect of Nef in primary T cells is unrelated to SERINC-antagonism (46), even though the virion-infectivity enhancing effect of Nef reportedly is (34).

Given these conflicting results, we aimed to further test the hypothesis that Nef enhances HIV-1 replication in a SERINC-dependent manner. To do this, we returned to the CD4-positive T cell line in which we originally observed a stimulation of growth-rate by Nef, an effect that was associated with Nef-mediated enhancement of virion-infectivity (CEM) (23). Here, we show that neither the attenuated replication-rate of *nef*-deficient HIV-1 in CEM T cells nor the reduced virion-infectivity of *nef*-deficient HIV-1 produced by these cells is rescued by CRISPR/cas9 editing of *serinc5*, either with or without additional editing of *serinc3*. These results support those documented using MOLT-3 and primary CD4-positive T cells and suggest that how Nef stimulates HIV-1 replication and virion infectivity remains unclear (46).

## Materials and Methods

### Cell Lines and Plasmids

HEK293T (a generous gift from Dr. Ned Landau) and HeLa TZM-bl cells (Dr. John Kappes and Xiaoyun Wu,: NIH AIDS Reagent Program, Division of AIDS, NIAID, NIH) (47, 48) were grown in DMEM media (Thermo Fisher Scientific) supplemented with 10% FBS (Hyclone) and 1% Penicillin/Streptomycin (Thermo Fisher Scientific). HeLa P4.R5 (obtained from Dr. Ned Landau) were cultured in 10% FBS, 1% Penicillin/Streptomycin and 1 µg puromycin. Both HeLa cell line derivatives express CD4, CXCR4 and CCR5 and contain either a Tat-inducible β-galactosidase (HeLa P4.R5) or both the β-galactosidase and luciferase (HeLa TZM-bl) genes under the transcriptional control of the HIV-1 LTR. CEM (a generous gift from Dr. Douglas Richman) and JTAg cells expressing (JTAg WT) or lacking SERINC3 and SERINC5 (JTAg SERINC3/SERINC5 KO; kindly provided by Dr. Heinrich Gottlinger) are T cell leukemic clones that were cultured in RPMI 1640 media plus 10% FBS and 1% Penicillin/Streptomycin (Thermo Fisher Scientific). The proviral plasmids pNL4-3 and pNL4-3ΔNef have been previously described (23). LentiCRISPRv2 (Addgene; Catalog #: 52961) contains a single guide RNA (sgRNA) targeting Exon 2 of SERINC3 (5’ATAAATGAGGCGAGTCACCG-3’) and was a gift from Dr. Massimo Pizzato (34). The lentiviral packaging (pxRSV-Rev, pMDLg/pRRE) and envelope (pMD2.G) plasmids were kindly provided by Dr. Dan Gibbs. LentiCRISPR-GFP encodes GFP in place of puromycin (49). Five sgRNAs targeting either Exon 1 or Exon 2 of SERINC5 were designed using an online CRISPR tool (benchling.com) and cloned into LentiCRISPR-GFP using previously described methods (50, 51). The sgRNA sequences were as follows: sgRNA-SERINC5(1), 5’-ACA GCACTGAGCTGACATCG-3’ ; sgRNA-SERINC5(2), 5’GCACTGAGCTGACATCGC GG-3’ ; sgRNA-SERINC5(3), 5’-CTTCGTTCAAGTGTGAGCTG’3’ ; sgRNA-SERINC5(4), 5’-CATCATGATGTCAACAACCG-3’ ; sgRNA-SERINC5(5), 5’-TGAGGGACTGCCGAATCCTG-3’. Briefly, sgRNA oligos were designed to produce the same overhangs after *BsmBI* digestion (5’-*CACCG*(sgRNA Oligo #1)-3’ ; 3’-*C*(sgRNA Oligo #2)-*CAAA*-3’). The oligos were phosphorylated (T4 Polynucleotide Kinase; NEB) and annealed in a thermal cycler according to the following conditions: 37°C for 30 minutes; 95°C for 5 minutes with a ramp down to 25°C at 5°C/minute. Diluted oligos (1:200) were ligated (T4 ligase; NEB) into dephosphorylated (Fast AP; Fermentas) and *BsmBI* digested (Fast *BsmBI*; Fermentas) LentiCRISPR-GFP by overnight incubation at 16°C, followed by transformation into Stbl3 bacteria (Thermo Fisher Scientific). Plasmid DNA was isolated from overnight bacterial cultures and verified via Sanger sequencing. *Generation of stable cell lines using CRISPR-Cas9:* To produce 3^rd^ generation lentiviral stocks, HEK293T cells were transfected with a total of 22.5 µg total plasmid according to the following equimolar ratios: 10 µg LentiCRISPR transfer plasmid (empty or containing sgRNAs against either SERINC3 or SERINC5), 5.9 µg pMDLg/pRRE, 2.8 µg pxRSV-Rev and 3.8 µg pMD2.G. Forty-eight hours later, concentrated lentivirus-containing supernatant was harvested following low-speed centrifugation, filtration (0.45 µm), and mixture with Lenti-X concentrator (Takara Bio) according to the manufacturer’s instructions. Briefly, 1 volume Lenti-X concentrator was mixed with 3 volumes clarified supernatant. The mixture was incubated for 30 minutes at 4°C, centrifuged at 1,500 x g for 45 minutes at 4°C, resuspended in 1 ml complete DMEM media and immediately stored at −80°C in single-use aliquots (100 µL).

We used previously validated sgRNAs to edit SERINC3 (34), whereas each of the five sgRNAs targeting SERINC5 were screened and the sgRNA which caused the most efficient editing in bulk transduced cells was chosen (data not shown). To create CEM cells knocked out for SERINC3 (S3-KO), we spinoculated 1 × 10^6^ cells with 100 µL lentivirus (LentiCRISPRv2-SERINC3 ; (34)) at 1,200 x g for 2 hours at 25°C. Puromycin (1 µg/ml) was added to cell cultures 72 hours post transduction to select for positive clones. Two weeks post-selection, genomic DNA was isolated from mock or lentiCRISPRv2-SERINC3 transduced CEM cells using the DNeasy Blood and Tissue Kit (Qiagen) according to the manufacturer’s instructions. Genome editing was assessed by Tracking of Indels by Decomposition (TIDE; (52)). PCR amplicons encompassing exon 2 of SERINC3 were produced using Taq 2x Master Mix (NEB) and the following primers: 5’-CAAATTACAACCAACTTGATTAACAACGACG-3’ and 5’-CTATAAAGCCTGATTTGCCTCGCTTTCTCTTC-3’. Clonal cell lines were isolated from bulk edited cultures using single-cell dilutions in a 96-well plate, followed by genomic DNA isolation and PCR amplification. Genome editing was verified via TIDE analysis.

To generate either single knockout or double-knockout cells lacking SERINC5, CEM WT or CEM S3-KO cells were spinoculated as described above with either empty lentivirus (LentiCRISPR-GFP) or virus containing sgRNA-SERINC5(4). Isolation of genomic DNA, PCR amplification and TIDE analysis were carried out seventy-two hours post transduction in a similar manner to the generation of S3-KO cells. We then expanded cells, which yielded two single SERINC5 knockout (S5-KO) clonal lines, and one SERINC3/SERINC5 knockout clonal cell line. We named these lines: SERINC5 knockout clone 8 (S5-KO (8)), SERINC5 knockout clone 11 (S5-KO (11)) and SERINC3/SERINC5 knockout clone 9 (S3/S5-KO (9)). The following primers were used to generate PCR amplicons for TIDE analysis: 5’-AGTGCCTGGCCATGTTTCTT-3’ and 5’-CATAGAGCAGGCTTCAGGAA-3’.

### HIV-1 production and titer

To produce replication-competent viruses, HEK293T cells (4 × 10^6^ /10 cm plate) were transfected with 24 µg of an infectious molecular clone of HIV-1 (NL4-3) or a mutant lacking the *nef* gene (NL4-3ΔNef) using Lipofectamine 2000 reagent according to the manufacturer’s instructions (Thermo Fisher Scientific). Virus-containing supernatants were collected forty-eight hours post transfection, clarified by low-speed centrifugation, and stored at −80°C. Viral titers were measured by infecting HeLa P4.R5 cells with diluted viral stocks in duplicate in a 48-well format for 48 hours. The cells were then fixed (1% formaldehyde; 0.2% glutaraldehyde) for 5 minutes at room-temperature, followed by overnight staining at 37°C with a solution composed of 4 mM potassium ferrocyanide, 4mM potassium ferricyanide, 2 mM MgCl_2_ and 0.4 mg/ml X-gal (5-bromo-4-chloro-3-indolyl-β-D-galactopyranoside). Infected cells (expressed as infectious centers (IC)) were quantified using a computer image-based method (53). (ABL Bioscience).

### HIV-1 infection and replication

To conduct HIV-1 replication studies, 1 × 10^6^ CEM cells (wildtype (WT); SERINC5 knockout clone 8 (S5KO (8)); SERINC5 knockout clone 11 (S5KO (11)); SERINC3/SERINC5 knockout clone 9 (S3/S5 K.O. (9))) were infected with NL4-3 (hereafter termed Nef+) or NL4-3ΔNef (hereafter termed Nef-) at a multiplicity of infection (MOI) of 0.01 overnight at 37°C in a 24-well plate format. The cells were then washed 3 times with 1 ml PBS (Corning), resuspended in 4 ml complete growth media (RPMI 1640), transferred to T25 labeled flasks and incubated at 37°C in an “upright” position for the duration of the experiment. Every three days, cultures were split 1:4 (1 ml cells; 3 ml media) and an aliquot (1 ml viral supernatant) stored at −80°C for quantification of HIV-1 replication (p24 antigen) by ELISA.

### Measurement of HIV-1 infectivity

Viral infectivity was quantified from virions produced in CEM cells infected with NL4-3 or NL4-3ΔNef at day 12 post-infection. A 20% sucrose cushion was used to concentrate virions via centrifugation according to the following parameters: 23,500 x g; 1 hour at 4°C. Viral pellets were resuspended in culture medium and dilutions used to infect 1.25 × 10^4^ HeLa TZM-bl cells in triplicate in a 96-well format. Forty-eight hours later, the culture medium was removed, and the cells were lysed using a luciferase reporter gene assay reagent (Britelite, Perkin Elmer). Infectivity (luciferase activity) was measured using a luminometer with data expressed as relative light units (RLU). These values were normalized to the p24 concentration of each sample and shown as RLU/p24.

### RT-qPCR and analysis

Total cellular RNA was extracted from 1 × 10^6^ JTAg WT, JTAg S3/S5 KO, CEM WT, CEM S5KO (8), CEM S5KO (11), and CEM S3/S5 KO cells using a Quick-RNA miniprep kit (Zymo Research), followed by treatment with RNase-free DNAse I (Zymo Research). Complementary DNA (cDNA) was generated from 250 ng of all extracted RNA samples using M-MLV RT (Thermo Fisher Scientific) and treated with RNaseOUT (Thermo Fisher Scientific). cDNA was mixed with the respective primer pairs and SyGreen Blue Mix (PCR Biosystems) following the manufacturer’s protocol in biological triplicate and performed using a LightCycler 96 real-time PCR machine (Roche). Quantification cycle values were normalized to a reference gene (*GAPDH*) and relative *SERINC5* or *SERINC3* gene expression ratios were calculated using the 2^-ΔΔCt^ method (54). The following primers were used for analysis: *SERINC5:* 5’-ATCGAGTTCTGACGCTCTGC-3’ and 5’-GCTCTTCAGTGTCCTCTCCAC-3’; *SERINC3* 5’-AATTCAGGAACACCAGCCTC-3’ and 5’-GGTTGGGATTGCAGGAACGA-3’; *GAPDH* 5’-TGCACCACCAACTGCTTAGC-3’ and 5’-GGCATGGACTGTGGTCATGAG-3’.

### Data analysis and presentation

Datasets were analyzed and combined in Microsoft Excel and GraphPad Prism 8.0 software. Where indicated, two-tailed unpaired or paired t-tests were performed. We utilized Adobe Photoshop and Illustrator CS6 for figure production.

## Results

### Generation of CEM cells lacking SERINC3 and SERINC5

We utilized CRISPR/Cas9 gene editing to determine whether the modulation of SERINC3 and SERINC5 is required for Nef to enhance HIV-1 replication in CEM cells. We chose the CEM cell line due to its ability to support robust Nef-dependent HIV-1 replication and virion-infectivity (23). We hypothesized that Nef would remain an important factor in promoting viral spread whether or not CEM cells expressed SERINC3 and SERINC5, a result in conflict with data obtained using Jurkat T cells but consistent with data recently obtained using MOLT-3 T cells. To test this, we created clonal cell lines lacking either SERINC5 or both SERINC3 and SERINC5 (described in Methods). We identified three clones containing indels within the SERINC5 target region: “clone 8” (S5-KO (8)), containing 4 and 13 base pair deletions, “clone 11”, containing 2, 11, and 13 base pair deletions (S5/KO (11)) and “clone 9”, containing 10 and 13 base pair deletions (S3/S5KO (9)) ; Figure 1A). SERINC3 knockout cells consisted of indels bearing a single base pair insertion and a single base pair deletion (S3/S5 KO (9); Figure 1C).

**Figure 1:**
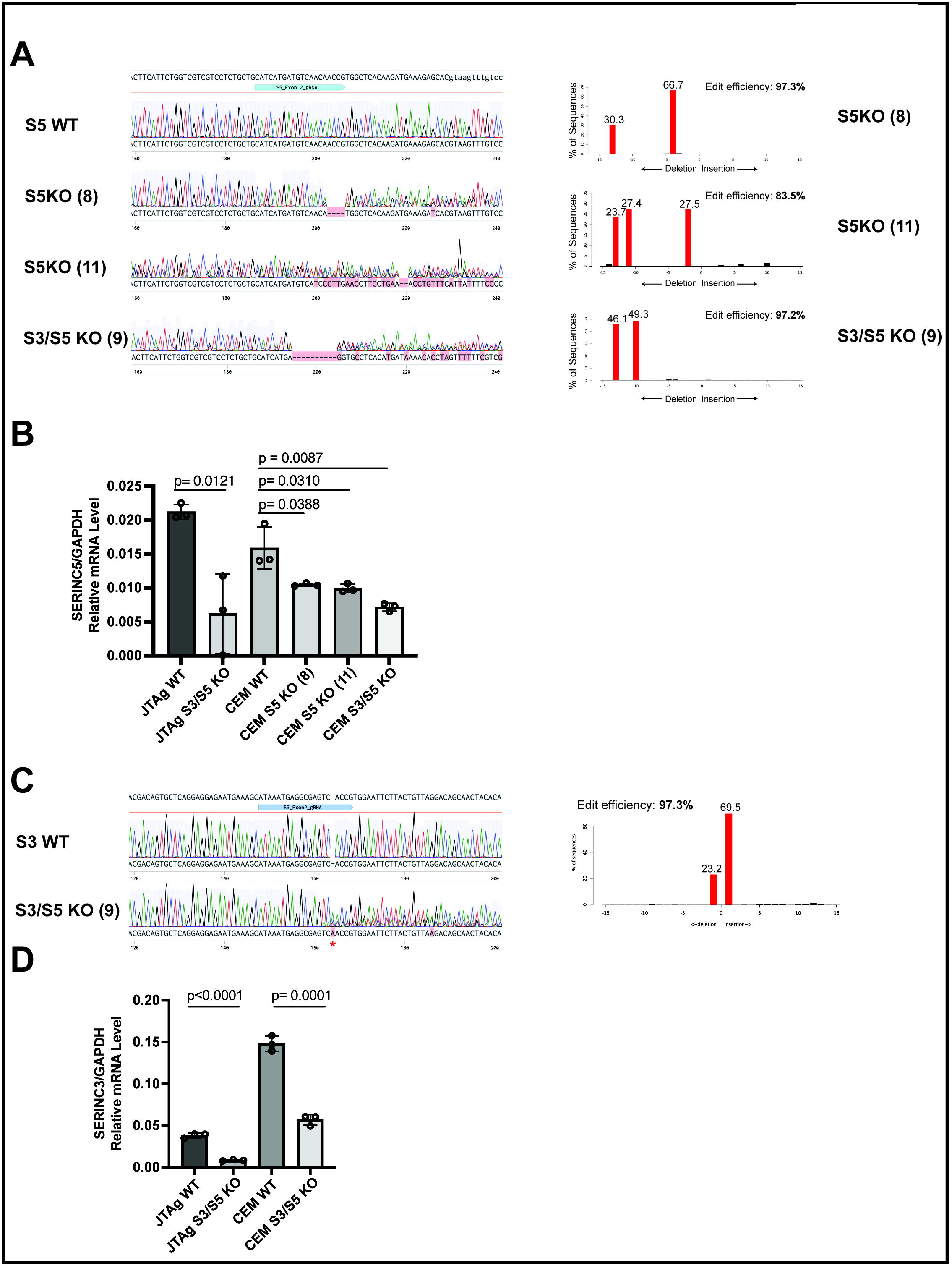
CRISPR/Cas9 editing of SERINC3 and SERINC5 in CEM cells. A.) *Left:* Chromatograms depicting a portion of exon 2 from SERINC5 wildtype (WT) or three SERINC5 knockout clones: clone 8 (S5KO (8)), clone 11 (S5KO (11)) or clone 9 (S3/S5 KO (9)). The guide RNA (gRNA) target site is shown as a blue arrow/annotation. Inserted nucleotides are highlighted red. Deletions are depicted as dashed line(s). *Right:* Tracking of Indels by Decomposition (TIDE) analysis of S5 KO clones. S5 KO (8) had 4 and 13bp deletions, S5KO (11) had 2,11 and 13 bp deletions and S3/S5 KO (9) had 10 and 13bp deletions. B.) Quantitative PCR (qPCR) showing the ratio of SERINC5 to GAPDH mRNA in JTAg and CEM cells expressing or lacking SERINC5. Results depict two independent experiments performed in triplicate. C.) *Left:* Chromatograms depicting a portion of exon 2 from SERINC3 wildtype (WT) or SERINC3/SERINC5 knockout clone 9 (S3/S5 KO (9)). The guide RNA (gRNA) target site is shown as a blue arrow/annotation. Inserted nucleotides are highlighted red. *Right:* TIDE analysis of S5 KO (9) predicted a 1bp insertion and 1bp deletion. D.) Quantitative PCR (qPCR) showing the ratio of SERINC3 to GAPDH mRNA in JTAg and CEM cells expressing or lacking SERINC3. Results depict two independent experiments performed in triplicate. Two-tailed unpaired t-tests were performed where indicated.

We attempted to validate editing in these cell lines using a monoclonal antibody targeting the extracellular domain of SERINC (55). However, this antibody did not detect endogenous SERINC5 in either Jurkat or CEM cells (data not shown). Instead, we reasoned that CRISPR/Cas9 editing of *serinc3* and *serinc5* would lead to a quantifiable decrease in mRNA transcripts due to nonsense mediated decay (NMD; (56)). To test this, we isolated RNA from CEM WT, CEM S5-KO (8), CEM S5-KO (11), CEM S3/S5-KO (9) and, as controls, JTAg WT and JTAg S3/S5-KO cells. In both JTAg and CEM, knockout clones expressed less *serinc5* (Figure 1B) or *serinc3* (Figure 1D) compared to wildtype clones. Given these results, these CEM cell lines served as the basis for our viral replication and infectivity studies.

### Optimal HIV-1 spread and viral infectivity in CEM cells is dependent on Nef but independent of SERINC5

To test whether Nef is required to counteract SERINC5 to enhance HIV-1 replication, we infected CEM wildtype (WT), S5-KO (8) and S5-KO (11) cells with either Nef expressing (hereafter termed Nef+: NL4-3) or Nef lacking (hereafter termed Nef-: NL4-3ΔNef) HIV-1 viruses at a multiplicity of infection (MOI)=0.01 infectious units per cell. The inocula for these growth-rate experiments were produced from HEK293T cells transfected with proviral plasmids, and the MOI was based on the infectivity of the virus stocks measured as infectious centers in cultures of CD4-positive HeLa-P4.R5 indicator cells (53). To infect 1×10^6^ CEM cells, we used amounts of Nef+ or Nef-virus stocks that yielded 10,000 infectious centers in the HeLa indicator assay. Although the infectivity of the viruses to CEM cells might be different than to CD4-HeLa cells, we chose this approach rather than normalizing the inocula to the content of p24 capsid antigen to adjust for the reduced infectivity of Nef-virions produced by the HEK293T cells (data not shown). The CEM cell cultures were split every 3 days and viral replication quantified by the amount of viral capsid (p24) within the supernatant (Figure 2A) measured by ELISA. The Nef+ viruses propagated more rapidly than the Nef-viruses, accumulating around 8.5-fold more p24 antigen in the culture supernates at 12 days post-infection in CEM WT cells (Figure 2B: Left panel). This corroborated the importance of Nef in enhancing viral replication in this *in vitro* system. If Nef-mediated modulation of SERINC5 were important for this phenotype, then the attenuated replication of Nef-virus should be “rescued” in cells lacking SERINC5. Instead, Nef enhanced HIV-1 replication in the CEM S5KO (8) and CEM S5KO (11) cell lines (Figure 2B: middle and right panels).

**Figure 2:**
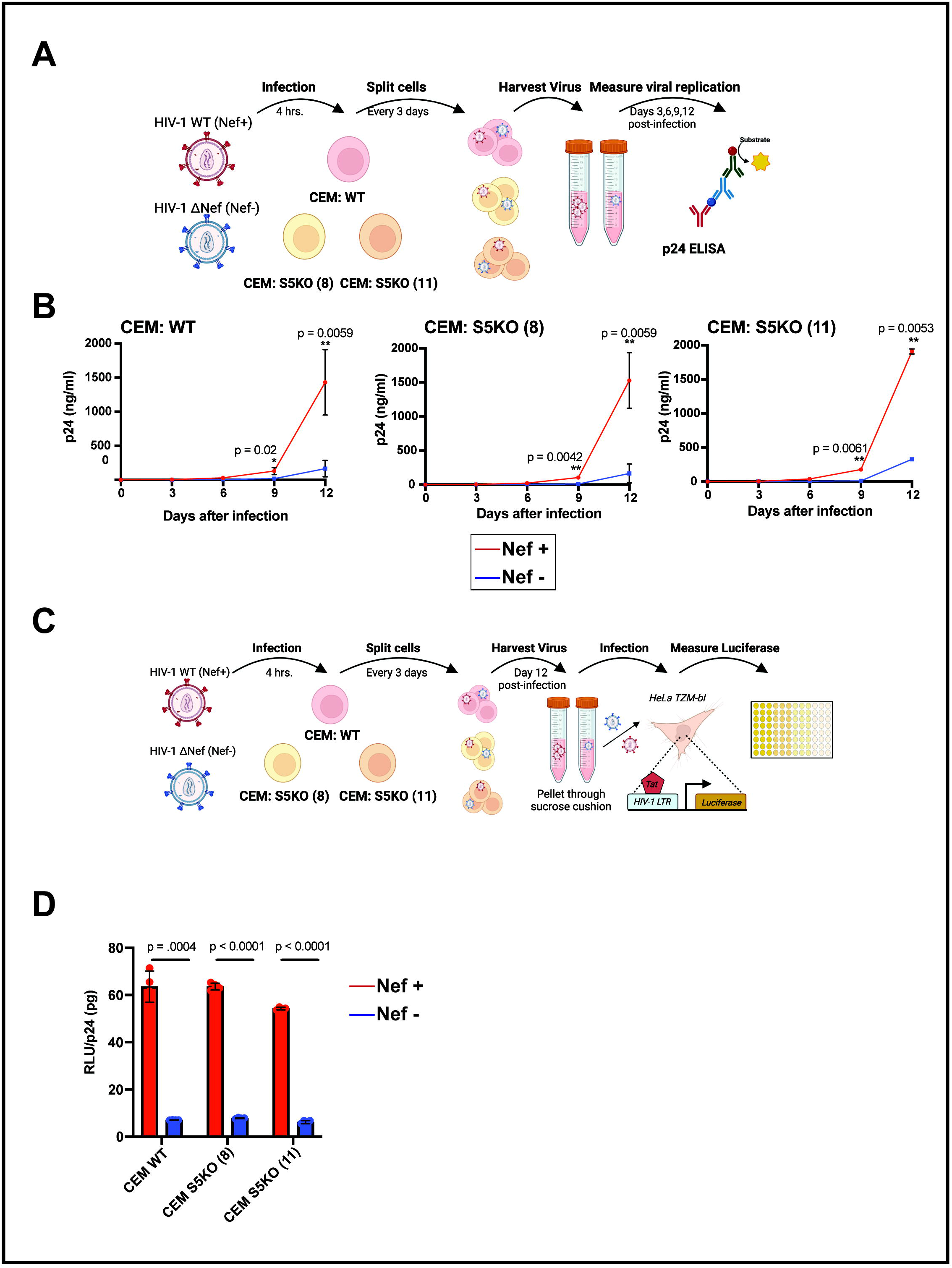
Nef enhances HIV-1 replication independently of SERINC5 in CEM cells. A.) Schematic of viral replication studies performed in CEM wildtype (WT), SERINC5 knockout clone 8 (S5KO (8)) or SERINC5 knockout clone 11 (S5KO (11)) cells. Each cell line was infected with either NL4-3 (termed Nef+) or HIV-1ΔNef (termed Nef-) at an MOI of 0.01. The cultures were split every 3-4 days and viral growth measured as indicated. Created with BioRender.com. B.) Viral replication quantified by p24 Capsid ELISA in the supernatants of CEM WT, S5KO (8) and S5KO (11) cultures. Results depict either two (WT, S5KO (8)) or one (S5KO (11)) independent infection measured in duplicate at each time point. C.) Schematic depicting measurement of single-cycle infectivity with virions produced from infected CEM WT, S5KO (8) or S5KO (11) cultures. HeLa TZM-bl indicator cells contain a luciferase gene under the transcriptional control of the HIV-1 LTR. Created with BioRender.com. D.) Infectivity data (relative luciferase units (RLU) normalized to p24 (RLU/p24)) from virions collected at day 12 post-infection from cultures of either CEM WT or S5KO (8) and S5KO (11) cells. The virions were partially purified by centrifugation through a 20% sucrose cushion before measuring infectivity (RLU in the HeLa TZM-bl assay) and p24 concentration (ELISA). Results are representative of one experiment performed in quadruplicate. Two-tailed paired t-tests were performed where indicated.

We next asked whether counteraction of SERINCs is necessary for Nef to enhance the infectiousness of virions produced by CEM cells, since Nef enhances virion-infectivity in a SERINC- and cell-type dependent manner (34). To test this, we collected virions from infected CEM cells at day 12 post-infection and measured single-cycle infectivity using HeLa TZM-bl reporter cells, which express luciferase under the transcriptional control of the HIV-1 LTR (Figure 2C). We used these cells because the luciferase read-out is more sensitive that the infectious center read-out of the HeLa-P4.R5 cells, and the concentration of Nef-virus produced by the CEM cultures was relatively low. Virions produced at day 12 post infection by Nef+ virus were around 8-fold more infectious per amount of p24 capsid antigen than those produced by Nef-virus, and this trend was observed regardless of SERINC5 expression (Figure 2D). Taken together, these data indicate that modulation of SERINC5 is not the primary mechanism by which Nef increases viral spread or infectivity in CEM cells.

### Nef enhances HIV-1 spread and virion-infectivity independently of SERINC3 in CEM cells

Whereas CEM cells express slightly less *serinc5* RNA than Jurkats, they express markedly more *serinc3* RNA (Figure 1, B and D). Thus, we sought to determine whether Nef-mediated modulation of SERINC3 influences viral growth rate and infectivity in these cells and might explain the *nef*-phenotype. We infected CEM WT and CEM cells lacking both SERINC3 and SERINC5 (S3/SKO (9)) with Nef+ and Nef – viruses and measured viral replication and infectivity in a similar manner as shown in Figure 2A and 2C. The absence of SERINC3 in CEM cells did not “rescue” Nef-virus in terms of either viral replication rate (Figure 3A, compare left and right panels) or virion infectivity (Figure 3B). Altogether, this data indicates that Nef increases virion infectivity as well as growth-rate in CEM T cells independently of SERINC3 and SERINC5.

**Figure 3:**
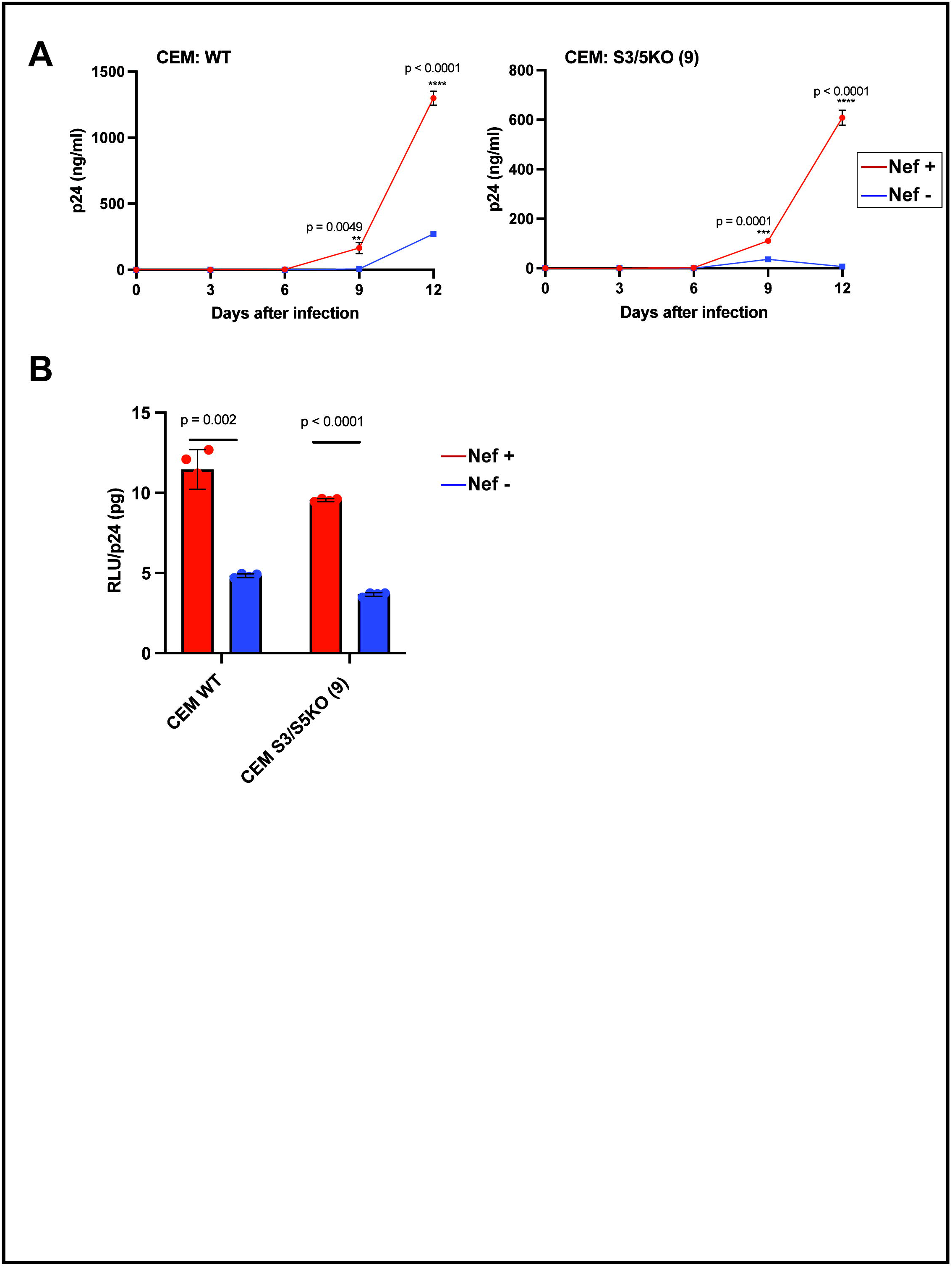
Nef enhances HIV-1 replication independently of SERINC3 in CEM cells. A.) Viral replication quantified by p24 Capsid ELISA in the supernatants of CEM WT, S5KO (8) and S5KO (11) cultures. Results depict one independent infection measured in quadruplicate at each time point. B.) Infectivity data (relative luciferase units (RLU) normalized to p24 (RLU/p24)) from virions collected at Day 12 post-infection from cultures of either CEM WT or S3/S5KO (9) cells. Results are representative of one experiment performed in quadruplicate. Two-tailed paired t-tests were performed where indicated.

## Discussion

In this study, we sought to determine whether Nef’s ability to enhance viral replication and/or infectivity in CEM T cells is primarily linked to modulation of SERINC3 and/or SERINC5. We present evidence that argues against this by showing that Nef increases viral replication rate in CEM cells lacking SERINC3 and SERINC5 and that virions of Nef+ virus are more infectious than those of Nef-virus regardless of whether SERINC3 and SERINC5 are expressed in the CEM cells that produced them.

Our finding that Nef enhances virion-infectivity independently of SERINCs in CEM T cells contrasts with reports in Jurkat and primary CD4-positive cells, where Nef mediated enhancement of infectivity appears to correlate with SERINC5 expression (34, 35). On the other hand, our observations herein using CEM cells are similar to recent results reported using MOLT-3 cells (46). In MOLT-3 cells, a chimeric HIV-1 virus bearing the SERINC5 antagonist (glycoMA) failed to substitute for Nef in rescuing HIV-1 infectivity (a property glycoMA should have if the sole function of Nef were to counteract SERINC5), and knockout of SERINC5 did not rescue the reduced growth rate of Nef-virus (46). Together, these data suggest that the SERINC-dependence of the Nef infectivity phenotype is cell-type dependent (34, 35, 46), with Jurkat being SERINC-dependent and CEM and MOLT-3 cells being SERINC-independent. Comparing virions produced from infected primary CD4-positive T cells either lacking or expressing SERINC5 seems essential to adjudicate the relevance of observations made using these T cell lines.

What mechanisms might explain Nef-mediated enhancement of infectivity and/or viral replication independently of SERINCs? One potential explanation could be Nef-mediated modulation of Src family kinases (SFKs). Nef binds several SFK members such as Lck, Hck, Lyn, and c-Src through a conserved proline-rich (PxxP) motif contained within Nef’s

Src homology region 3 (SH3) binding domain (57, 58), and primary Nef isolates from HIV-1 Group M display a conserved ability to activate SFK’s (59). Nef mutants lacking the PxxP motif were reportedly unable to enhance HIV-1 replication in peripheral blood mononuclear cells (58). However, other studies reported that mutation of Nef’s PxxP motif yielded little to no difference in HIV-1 replication within MOLT-3 or primary CD4-positive T cells, and in CEM T cells the attenuated phenotype of mutants of the SH3 binding domain was modest and seemed attributable to reduced expression of Nef (25, 46, 60). Analyzing HIV-1 replication in T cells lacking one or more SFKs may be necessary to adequately assess whether Nef’s ability to enhance HIV-1 replication is mediated by these interactions.

One notable similarity between MOLT-3 and CEM lymphoblastoid cells that distinguishes them from Jurkat cells is that they do not express the T cell receptor (TCR) at their surfaces (61, 62). Given that Nef reportedly mimics TCR-signaling (22, 63), another possibility is that MOLT-3 and CEM cells reveal a SERINC-independent growth-rate Nef phenotype that is exaggerated by the absence of constitutive TCR signaling in these cells. This could be consistent with the initial observations that the activation-state of primary CD4-positive T cells affects the Nef growth-rate phenotype (24). Nonetheless, how TCR signaling would affect virion-infectivity is obscure.

Lastly, the explanation might reside in Nef’s ability to interact with components of clathrin-mediated trafficking pathways and to modulate many cellular membrane proteins, one of which might be currently unidentified but underlie the infectivity phenotype in MOLT-3 and CEM cells. Nef binds AP complexes via a conserved sorting signal near its C-terminus: 160ExxxLL164,165 (64). This “di-leucine motif” is required for both Nef-mediated CD4 downregulation and optimal viral infectivity in CEM cells (65). The Nef LL164/165AA mutant, which is unable to bind AP-2 (66), replicates poorly in MOLT-3 cells (46). Residues located within Nef’s core domain and required for downregulating CD4 and interacting with Dynamin-2, a “pinchase” of clathrin coated pits, are also required for enhancement of viral replication in MOLT-3 cells (46, 67, 68). Finally, Nef enhances HIV-1 replication in the absence of CD4 downregulation in MOLT-3 cells (46) and in CEM-derived cells (A2.01) engineered to express a CD4 lacking a cytoplasmic domain that is unresponsive to Nef (69). These observations regarding CD4 are critically important, since CD4 is itself a potent inhibitor of infectivity that is counteracted by Nef (11, 12). In the CEM experiments herein, CD4 downregulation by Nef could conceivably account for the observed infectivity and growth-rate phenotypes. Nonetheless, even if that were the case, the data would indicate that the contribution of CD4 to these phenotypes far outweighs the contribution of SERINC3 and SERINC5 in these cells.

Our study has several caveats. As noted above, CEM cells, like MOLT3 cells, might not reflect the role of Nef in primary T cells or macrophages. Also, quantitative experimental variation is evident in the CEM system, with the *nef* virion-infectivity phenotype varying between 2- and 8-fold (compare Figures 2 and 3). Nonetheless, a *nef*-phenotype was always apparent in these cells, and it was unaffected by knock-out of *serinc5* and *serinc3*.

Overall, whether Nef provides a direct, positive effect on viral replication or instead is counteracting a still unidentified restriction factor remains to be determined. Utilizing both MOLT-3 and CEM cells might facilitate answering this question, providing a more complete understanding of the enigmatic yet important virologic effects of Nef.

## Author Contributions

**Peter W. Ramirez:** Conceptualization, Methodology, Investigation, Writing – original draft & review and editing **Francisco M. Acosta, Marissa Suarez, Aaron A. Angerstein, Thomas Vollbrecht**: Validation, Methodology, Investigation, Writing – review and editing **Jared Wallace, Ryan M. O’ Connell**: Resources **John Guatelli:** Conceptualization, Methodology, Writing – review and editing.

## Acknowledgments

We thank Dr. Vicente Planelles for the LentiCRISPR-GFP plasmid. We thank The Pendleton Charitable Trust for equipment. P.W.R. was partly supported by NIH grant K12GM068524. This work was supported by NIH grant R01 AI129706 to J.G.

